# Lateralization of orthographic processing in fixed-gaze and natural reading conditions

**DOI:** 10.1101/2022.06.20.496859

**Authors:** Ádám Nárai, Zsuzsanna Nemecz, Zoltán Vidnyánszky, Béla Weiss

**Author notes:** **Corresponding author:** Béla Weiss, Brain Imaging Centre, Research Centre for Natural Sciences, Magyar tudósok körútja 2. 1117 Budapest, Hungary. The first two authors contributed equally to this work.

## Abstract

Lateralized processing of orthographic information is a hallmark of proficient reading. However, how this finding obtained for fixed-gaze processing of orthographic stimuli translates to ecologically valid reading conditions remained to be clarified. To address this shortcoming, here we assessed the lateralization of early orthographic processing in fixed-gaze and natural reading conditions using concomitant eye-tracking and EEG data recorded from young adults without reading difficulties. Sensor-space analyses confirmed the well-known left-lateralized negative-going deflection of fixed-gaze EEG activity throughout the period of early orthographic processing. At the same time, fixation-related EEG activity exhibited left-lateralized followed by right-lateralized processing of text stimuli during natural reading. A strong positive relationship was found between the early leftward lateralization in fixed-gaze and natural reading conditions. Using source-space analyses, early left-lateralized brain activity was obtained in lateraloccipital and posterior ventral occipito-temporal cortices reflecting letter-level processing in both conditions. In addition, in the same time interval, left-lateralized source activity was found also in premotor and parietal brain regions during natural reading. While brain activity remained left-lateralized in later stages representing word-level processing in posterior and middle ventral temporal regions in the fixed-gaze condition, fixation-related source activity became stronger in the right hemisphere in medial and more anterior ventral temporal brain regions indicating higher-level processing of orthographic information. Although our results show a strong positive relationship between the lateralization of letter-level processing in the two conditions and suggest lateralized brain activity as a general marker for processing of orthographic information, they also clearly indicate the need for reading research in ecologically valid conditions to identify the neural basis of visuospatial attentional, oculomotor and higher-level processes specific for natural reading.

## 1 Introduction

Lateralization of orthographic processing is considered to be a marker of fluent reading, and its impairment was shown to be an indicator of reading difficulties. Accordingly, identification of neural mechanisms that subserve lateralized processing of orthographic information would enhance the understanding of reading processes and could also contribute to the development of more efficient training programs aimed for mitigation of reading difficulties. However, despite its significant role in functioning and development of modern societies, the investigation of brain mechanisms of reading has been limited to fixed-gaze experimental conditions for multiple decades.

Previous research investigating orthographic processing in fixed-gaze conditions revealed left-lateralized negative deflection of electroencephalogram (EEG) activity at the latency of the first negative (N1 or N170) component (Bentin et al., 1999; Rossion et al., 2003) and also indicated more negative-going event-related potentials (ERPs) for words compared to consonant strings during the second negative and third positive components in left occipito-temporal electrodes (Cohen et al., 2000). Studies applying magnetoencephalography (MEG), functional magnetic resonance imaging (fMRI) and invasive recordings identified the brain sources presumably contributing to the lateralization of these ERP components that most likely reflect subsequent stages of letter-level, word-level and higher-level processing of orthographic information (Halgren et al., 1994, 2006; Lerma-Usabiaga et al., 2018; Thesen et al., 2012). The N1 ERP component is assumed to indicate letter-level processing in extrastriate occipital (Barzegaran & Norcia, 2020; Boros et al., 2016; Gold & Rastle, 2007; Hervais-Adelman et al., 2019; Posner et al., 1988; Puce et al., 1996; Temple et al., 2001; Woolnough et al., 2020) and left ventral posterior occipito-temporal cortices forming the letter-form area (Lerma-Usabiaga et al., 2018; Thesen et al., 2012). After the occipito-teporal N1 ERP component word-level processing emerges and exhibits a source activity peak at around 220 ms (Thesen et al., 2012) in the visual word-form area (VWFA), a brain region located anterior to the letter-form area (Cohen et al., 2000; Dehaene et al., 2002, 2005; Dehaene & Cohen, 2011; McCandliss et al., 2003; Price & Devlin, 2011; Thesen et al., 2012).

Hemispheric lateralization of orthographic processing has been shown to change with age (Spironelli & Angrilli, 2009), especially during childhood, when reading skills are under active development. The sensitivity of the N1 ERP component to words was found to follow an inverted “U” shape from kindergarten to 5^th^ grade (Maurer et al., 2011), and in line with this inverted “U” model of visual expertise development, several studies investigating 9-12 year-old children found rightward instead of leftward lateralization of the word-evoked N1 ERP component (Fraga González et al., 2014, 2016; Setten et al., 2019). Studies based on fixed-gaze experimental paradigms also revealed enhanced brain activation in the left occipito-temporal region for processing of orthographic information in experienced readers compared to beginners, the dominance of the left hemisphere in orthographic processing was shown to become more prominent with the advancement of reading skills (Brem et al., 2010; Dehaene et al., 2010, 2015; Pegado et al., 2014). The pseudoword-evoked N1 ERP component was found to be more leftward lateralized as reading expertise increased (Pegado et al., 2014), similarly to the leftward lateralization of the N1 component evoked by familiar orthographic stimuli (Maurer et al., 2008) that increases with reading acquisition (Brem et al., 2010). Moreover, using fMRI, the left-lateralized activation in the VWFA was shown to increase with literacy (Dehaene et al., 2010, 2015). Besides their sensitivity to reading expertise, brain imaging markers of lateralized orthographic processing have also been shown to indicate reading difficulties. It has been proposed that compromised expertise-driven specialization of early orthographic processing to the left hemisphere might be closely related to poor reading skills in developmental dyslexia (Brambati et al., 2006; Helenius, 1999; Mahé et al., 2012, 2013; Paulesu et al., 2001; Paz-Alonso et al., 2018; Richlan et al., 2009, 2011). A reduced lateralization of the first negative event-related component was found for dyslexics compared to control participants (Fraga González et al., 2014, 2016; Helenius, 1999; Mahé et al., 2012, 2013).

In contrast to fixed-gaze processing of orthographic information, natural reading is an active sensory-motor process relying on consecutive reallocation of visuospatial covert attention (Kornrumpf et al., 2016) and saccadic eye movements (Grainger et al., 2016; Rayner, 1998, 2009; Yarbus, 1967). Accordingly, it is an essential question how the enormous body of experimental evidence collected on hemispheric specialization of fixed-gaze orthographic processing translates to ecologically valid reading conditions (Dimigen et al., 2011; Hauk & Weiss, 2020; Hutzler et al., 2007; Kornrumpf et al., 2016; S. C. Sereno et al., 1998; S. Sereno & Rayner, 2003; Temereanca et al., 2012; Weiss et al., 2016; Weiss, Nárai, et al., 2022). In our recent study, we assessed the lateralization of fixation-related EEG activity (FREA) during natural reading in dyslexic and control young adults (Weiss, Nárai, et al., 2022). We found that occipito-temporal FREA is lateralized to left and then to right posterior electrodes at the latency of the first negative FREA component and during later stages of orthographic processing, respectively. We also revealed that the early leftward lateralization of negative-going FREA deflection is specific for first fixations, it is most prominent at default inter-letter spacing and it deteriorates in participants with reading difficulties. However, to our knowledge, this is the only study on the lateralization of orthographic processing during natural reading, and thus, it remains to be clarified how these findings relate to the earlier results obtained in fixed-gaze conditions as well as what the source-space correlates of significant FREA lateralization revealed by sensor-space analyses are. To address this shortcoming, simultaneous recording of eye-tracking (ET) and high-resolution brain imaging data is required (Weiss et al., 2021).

In this study, we recorded concomitant ET and high-density EEG data in fixed-gaze and natural reading conditions. During the fixed-gaze paradigm, participants performed a one-back task using word, false-font and phase-randomized stimuli, while in the natural reading condition, isolated meaningful sentences were read silently in an ecologically valid manner. We tested the hemispheric specialization of orthographic processing by assessing the lateralization of fixed-gaze and fixation-related brain activity using sensor-space and source-space analyses of EEG recordings. While for statistical analysis of fixed-gaze EEG we used a traditional approach based on individual averages, in the case of EEG data recorded during natural reading, we controlled for potential confounding effects of eye movements by regressing the eye-movement covariates out from FREA at individual level. At group level, we tested the significance of brain activity lateralization in all experimental conditions, and also assessed the dependence of FREA and its lateralization on the rank of fixations. To evaluate the relationship between early orthographic processing in fixed-gaze and natural reading conditions, we performed correlation analysis between the lateralization of fixed-gaze and fixation-related EEG activity at the latency of first negative occipito-temporal peaks.

## 2 Material and methods

### 2.1 Participants

Thirty-nine young adults without reading difficulties participated in this study. One participant was excluded due to poor eye-tracking data quality. In the remaining group of 38 participants, 24 were female, average age was 20.76 years with 1.60 years standard deviation (SD). According to the laterality quotient (LQ) of the Edinburgh Handedness Inventory (Oldfield, 1971), there were 35 right handed, one left handed and two ambidextrous (LQ: 0 and 43) participants. Participants’ reading ability was assessed by a non-word reading test using the Hungarian adaptation (3DM-H; (Tóth et al., 2014)) of the Differential Diagnosis Dyslexia Maastricht Battery (3DM Battery; (Blomert & Vaessen, 2009)). In our recent study, we investigated the effects of reading ability on the lateralization of orthographic processing during natural reading (Weiss, Nárai, et al., 2022), and for validation of participants’ reading ability, we compared the behavioral and eye-movement measures of control readers to those of participants with official diagnosis of developmental dyslexia using hypothesis testing and machine learning approaches (Szalma et al., 2021; Szalma & Weiss, 2020; Weiss, Nárai, et al., 2022; Weiss, Szalma, et al., 2022). Here, to validate the reading ability of participants, we also contrasted their behavioral performance to the performance of control readers in our previous study (Weiss, Nárai, et al., 2022). Considering the non-word reading fluency score, the participants of the current study performed similarly or even better (Welch’s t-test: t=-2.27, P=0.027; Mann-Whitney U test: U=320, P=0.025) than the young adult control readers of our former study.

All participants were native speakers of Hungarian, unfamiliar with the Armenian alphabet, and had normal or corrected-to-normal vision. None of the participants had any history of neurological or psychiatric diseases. The study was approved by the Committee of Science and Research Ethics of the Medical Research Council, the ethical approval was issued by the Health Registration and Training Center (092026/2016/OTIG). All methods were carried out in accordance with the approved guidelines and subjects gave written informed consent before starting the experiment.

### 2.2 Stimuli and experimental procedures

In this study, two paradigms, a natural reading task and a fixed-gaze one-back task were performed in one session. The order of experimental paradigms was randomized across the subjects.

During the natural reading task, participants read isolated meaningful Hungarian sentences while their eye movements and electroencephalogram (EEG) activity were recorded. One hundred sentences were randomly chosen from the same pool of sentences that were also used in our recent study on lateralization of orthographic processing (Weiss, Nárai, et al., 2022). The sentences originated from the Hungarian Electronic Library (https://mek.oszk.hu), their order was randomized across the participants. Each sentence was 140-145 characters (20.24±0.39 words) long, started with a capital letter, contained only words with lemma and word frequencies larger than 0.01 per million and had neither special nor numeric characters except for ommas. Sentences were presented in four blocks using 13 pt Courier New font on a 23.5″ liquid-crystal display (LG Electronics Inc. 24M47VQ-P) with a resolution of 1920×1080 pixels and 60 Hz refresh rate. Viewing distance was 56 cm. Black letters were presented on a 53.3×30° (degrees of visual angle) white background with left and right horizontal margins of ∼5.8°. Average character width was ∼0.23°. All sentences consisted of a single text line aligned to left. Vertically centered fixation points (4 pixel diameter), placed ∼0.8° left and right from text boundaries were used for controlling sentence presentation and for eye-tracking quality validation. Each sentence trial started with only the left fixation point presented and participants were instructed to gaze on this point until it disappeared and the text line appeared with the right fixation point (Fig. 1a), then read the text carefully at their own pace and gaze at the right fixation point until the text automatically disappeared after the validation procedure (see the Supplementary Material, section 1.1). To assure that participants pay attention to the task, a test statement about the last read sentence was randomly shown after 25 % of sentences, and participants had to report by pressing a keyboard button whether the statement is true (key J) or false (key F).

**Figure 1.**
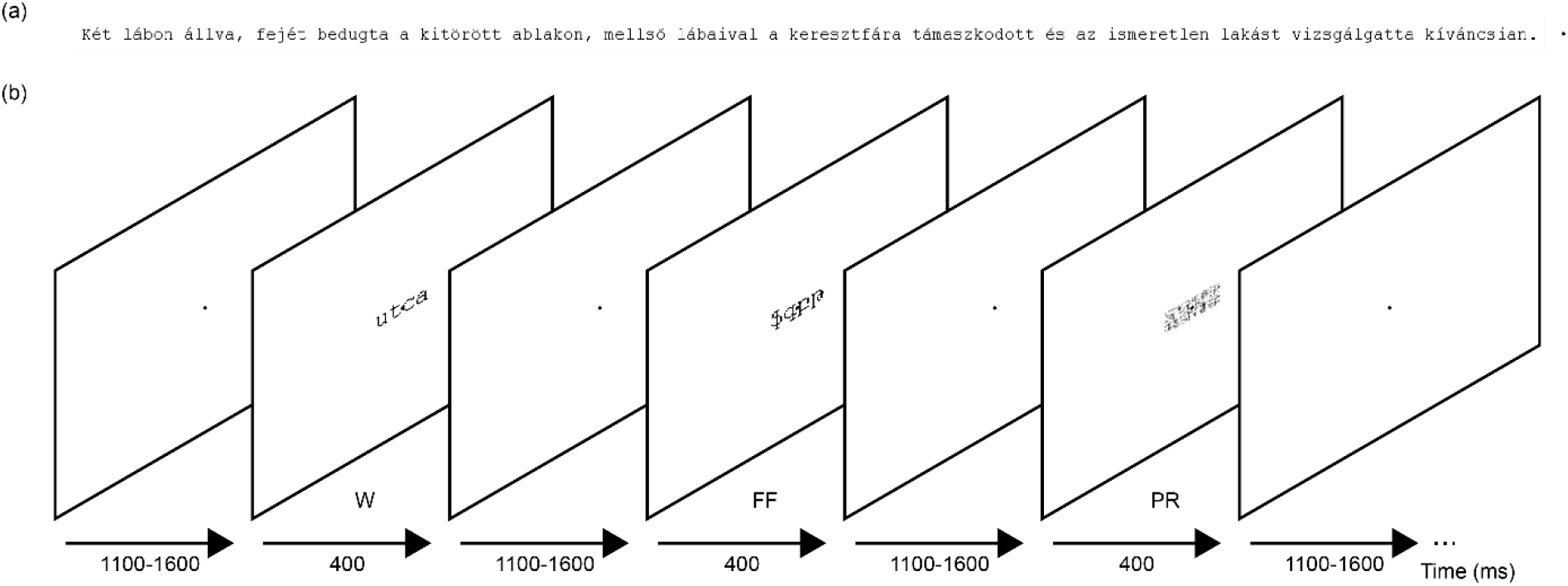
Sample visual stimuli. During the natural reading paradigm Hungarian sentences consisting of a single text line were presented along with a fixation point that was placed to the right of text stimuli (a). The schematic of the fixed-gaze one-back paradigm is shown in panel (b). Hungarian words (W), Armenian character strings representing false-font strings (FF) and phase-randomized Hungarian words (PR) were presented together with a centered fixation point for 400 ms in a pseudo-random order. The presentation of W, FF and PR stimuli was separated by a black fixation point centered on a white background. This fixation dot was presented for a randomized time interval between 1100 and 1600 ms resulting thus in a 1500-2000 ms stimulus onset asynchrony.

In our recent study, we revealed impaired lateralization of early orthographic processing during natural reading in dyslexic adults that suggests compromised letter-level processing of orthographic information in participants with reading difficulties (Weiss, Nárai, et al., 2022). Accordingly, in addition to investigating the relationship between the early lateralization of natural reading and fixed-gaze orthographic processing, the fixed-gaze paradigm was designed to also assess the effects of reading expertise on the lateralization of letter-level orthographic processing. The effects of letter familiarity were investigated by contrasting the lateralization correlates of processing real words (W, familiar letters of the Hungarian alphabet) and false-font stimuli (FF, strings of unfamiliar Armenian letters). Moreover, phase randomized stimuli (PR) were also included as a reference condition (Fig. 1b). A one-back task that could be carried out for all the three types of stimuli was used in the case of the fixed-gaze paradigm (Brem et al., 2018). One hundred sixty common Hungarian words were selected from the Hungarian National Corpus (Oravecz et al., 2014). Four- and five-letter long nouns and adjectives were used, 40 words per category, having word frequency between 20.25 and 1445 words per million. Armenian character strings were generated by replacing the letters of the Hungarian words by a random Armenian character. Seven Armenian characters closely resembling Hungarian alphanumeric characters `(‘զ’,’հ’,’շ’,’ո’,’ս’,’ց’,’օ’)` were excluded. Phase-randomized stimuli were generated by applying the weighted mean phase method (Dakin et al., 2002) on bitmaps of Hungarian words. A fixation point of a two-pixel diameter was presented in the screen center. Hungarian words and false-font strings were presented at screen center in black on a white background using 13 pt Courier New font. Phase-randomized stimuli derived from the four- and five-letter long Hungarian words were presented at the screen center in a 1.14°×0.44° and 1.42°×0.44° area, respectively. The W, FF and PR stimuli were presented for 400 ms in a pseudo random order. Consecutive repetition of the same condition for more than three times was not allowed. The inter-stimulus interval was sampled from a uniform distribution between 1100 and 1600 ms. Ten percent of stimuli, i.e. 16 stimuli of each condition were repeated immediately in a random fashion to create a one-back task. Participants were instructed to gaze at the fixation point and press a key when two consecutive stimuli were identical. They had to press the F key with their left or the J key with their right index finger. The keys used to collect the responses were randomized across the subjects. The performance was above 85 % on average in all the three conditions (W: 98.9%±3.7%; FF: 96.7%±3.7%; PR: 86.5%±8.2%) indicating that participants paid attention to the task. Trials with repeated stimuli were excluded from EEG analysis. A five-second-long break was pseudo randomly placed after 10 trials on average. During the short breaks participants had the opportunity to rest their eyes. The frequency of these breaks was increased upon participants’ request or if the experimenter found it necessary to reduce the number of eye blinks occurring during the presentation of W, FF and PR stimuli. Stimulus presentation was divided into nine blocks, and participants had the opportunity to rest between the blocks.

Generation and presentation of stimuli, control of experimental procedures and collection of subjects’ responses was performed using custom written scripts and the Psychophysics Toolbox 3 (v. 3.0.9) (Brainard, 1997; Kleiner et al., 2007; Pelli, 1997) under MATLAB R2008a (The MathWorks Inc., Natick, MA, USA).

### 2.3 Data acquisition

EEG data were acquired using a BrainAmp Standard amplifier with a 96-channel actiCAP active electrode system and the BrainVision Recorder 1.2 software (Brain Products GmbH, Munich, Germany). EEG was recorded using 95 electrodes placed according to the 10-5 international standard, and one electrode was put below the left eye for vertical electrooculography (vEOG). All channels were referenced to the right mastoid (TP10), while the ground electrode was at AFz. Electrode impedances were kept below 5 kΩ and data were sampled at 500 Hz. ET data were recorded from the left eye using an EyeLink 1000 Plus system (SR Research Ltd, Ottawa, Ontario, Canada) with 1000 Hz sampling rate. Calibration was carried out using the built-in randomized nine-point routine, squeezing the calibration area vertically during the natural reading task to relocate the fixation points closer to the horizontal midline. ET accuracy and precision was validated before the presentation and after reading of all text lines by using a 1°×1° square around the left and right fixation points presented at the beginning and the end of text stimuli, respectively (Supplementary Material, section 1.1). For synchronization of EEG and ET recordings, trigger signals and messages were sent to the EEG system through a standard trigger port and to the ET system using the Ethernet protocol. Trigger signals coded the appearance and disappearance of text lines. To validate the timing of trigger signals and messages, multiple tests were performed (Supplementary Material, section 1.2).

### 2.4 Processing of eye-tracking data

ET data recorded during the natural reading task was preprocessed in two stages. First, segments of horizontal and vertical gaze positions were extracted from between “text line appeared” and “text line disappeared” triggers. These segments were filtered and denoised using a Savitzky-Golay FIR filter as implemented in the adaptive algorithm toolbox (Nyström & Holmqvist, 2010). The end of the segments was corrected using this filtered data. The time at which the horizontal gaze position crossed the most-right black pixel of the text line for the first time was considered as the correct end of the reading segments. Raw ET data segments were trimmed at these line-end times. In the second stage, trimmed ET segments were processed with an adaptive algorithm (Nyström & Holmqvist, 2010) using the following settings: max. saccade velocity=1000°/s; max. saccade acceleration=100000°/s^2^; min. fixation duration=40 ms; min. saccade duration=10 ms; α_AA_=0.7; β_AA_=0.3; initial saccade peak velocity detection threshold PT1=50°/s. This procedure yielded the timing and characteristic features of saccades and fixations. ET data processing was performed in MATLAB R2016b (The MathWorks Inc., Natick, MA, USA) with custom scripts based on the adaptive algorithm toolbox provided by dr. Marcus Nyström.

### 2.5 Preprocessing of EEG data

Processing of the EEG data recorded during natural reading closely followed the data processing pipeline suggested in (Weiss et al., 2016) with some amendments. Preprocessing started with zero-phase digital filtering of EEG time-series, applying a 4th-order Butterworth band-pass (0.5– 70 Hz) filter and a 50 Hz notch filter (quality factor Q=45) using the filtfilt() MATLAB function. EEG data between “text line appeared” and “text line disappeared” triggers were segmented into 500 ms long epochs for semi-automatic artifact rejection. Automatic artifact detection was carried out using EEGLAB (Delorme & Makeig, 2004) functions with the following parameters: simple amplitude limit with [-100 100] μV thresholds; trend limit with a maximum absolute slope of 25 μV within a 500 ms epoch (R^2^≥0.5); spectral limit with [-50 50] dB threshold in the 0-2 Hz and [-100 25] dB in the 20-40 Hz frequency bands; improbable data kurtosis criterion with 5 SD threshold for both single-channel and global data. Since eye movements were an integral part of this paradigm, results of automatic artifact detection were visually inspected, and segments containing only eye blinks or saccadic artifacts were unmarked. This process was aided by blink detection applying the ET data (zero horizontal and vertical gaze positions were regarded as blinks) as well as using an EOG based method. For this purpose, the absolute maximum of vEOG-AF3 bipolar channel activity was thresholded within the 500 ms segments. The marked artifact contaminated segments were excluded from further analyses.

The remaining EEG segments were subjected to independent component analysis (ICA) using the runica() EEGLAB function with ‘extended’, 1, ‘stop’, 1e-7, ‘maxsteps’, 1024 parameters. Artifactual independent components (ICs) were detected using the ICLabel (Pion-Tonachini et al., 2019) and SASICA (Chaumon et al., 2015) toolboxes with default settings, except the SNR measure of SASICA. We defined the SNR measure as the ratio of average FREA between the [50 200] ms and [-40 0] ms time ranges with the intention of capturing saccade-related artifacts. Results were validated by visual inspection of ICs’ topography, spectral properties and single-trial fixation-related activation images.

Artifactual ICs were eliminated and the obtained ICA weights were applied to the continuous filtered EEG data. For sensor-space analysis, surface Laplacian also known as scalp current density (SCD) transformation was applied on cleaned EEG data using the CSD Toolbox (Kayser & Tenke, 2006) with unit sphere radius, m=4 (spline flexibility), λ=10^−5^ (smoothing constant) parameters and the maximum degree of Legendre polynomials set to 10. SCD transformed data can be considered reference independent, the transformation enhances local activity and suppresses activity with broader spatial extent (Perrin et al., 1987). Accordingly, the SCD transformation can also be used to reduce the effects of volume conduction and saccadic potentials (Babiloni et al., 1996; Melloni et al., 2009). As Laplace transformed EEG rather reflects distribution of scalp current density instead of scalp potentials, here we denote fixed-gaze stimulus-onset-related EEG activity and fixation-related EEG activity by the FGEA and FREA abbreviations, respectively, instead of using the terms event-related potential (ERP) and fixation-related potential (FRP). Furthermore, a 35 Hz 4^th^-order low-pass Butterworth filter was applied to remove unwanted high-frequency components. For source-space analysis, cleaned EEG data were average referenced. Both Laplace transformed and average referenced FREA trials were extracted from continuous EEG using a [-250 600] ms time window triggered to saccade end times obtained from the ET data. Saccade end times were considered as the onset of subsequent fixations. Trials overlapping with artifactual segments, the first 650 ms and last 200 ms of text line presentation were omitted from analyses. Finally, FREA trials were baseline corrected by subtracting the average activity of the [-50 0] ms time interval.

EEG data recorded during the fixed-gaze task were preprocessed by filtering and semi-automatic artifact detection steps with parameters identical to those applied for preprocessing of natural-reading EEG. However, in this case, artifact detection was directly performed on the EEG epochs that were used for statistical analysis. Epochs contaminated by saccadic eye movements were excluded from further analyses. Artifacts related to eye blinks and microsaccades and those present throughout the recording sessions were eliminated by applying the ICA weights obtained for the natural reading task. Laplace transformation and 35 Hz low-pass filtering was also applied for sensor-space analysis, while average referenced EEG was used for source estimation. The epochs were obtained by triggering the EEG data to the onset of stimuli using a [-250 600] ms time window. Finally, the obtained trials were baseline corrected by subtracting the average activity of the [-250 0] ms time interval.

EEG preprocessing was performed in MATLAB R2016b (The MathWorks Inc., Natick, MA, USA) using custom written scripts and the EEGLAB toolbox.

### 2.6 Estimation of brain source activity

Estimation of distributed source activity was based on average referenced fixed-gaze and fixation-related EEG trials. The potential confounding effects of eye-movement covariates were regressed out from average referenced sensor-space FREA similarly as for the statistical analyses performed in sensor space after Laplace transformation (section 2.7 and Supplementary Material, section 1.3).

A source space with icosahedron subdivisions and 3.1 mm spacing between grid points was created inside the FreeSurfer’s FSAverage template (Fischl, 2012). The forward solution was calculated using the boundary-element model. The noise covariance matrices were computed with automated model selection (Engemann & Gramfort, 2015). Two noise covariance matrices were computed for all participants, one for fixed-gaze and another for fixation-related EEG using the 250 ms and 50 ms long baseline data, respectively. Inverse operators were calculated by entering the matrices composed of the noise covariance matrices, the forward model, and the source covariance matrices to singular value decomposition. Dipole orientations were not fixed, but only limited to an orientation perpendicular to the cortex with a weight value of 0.2 for dipole components tangential to the cortical surface, and a depth weighting exponent of 0.8. The obtained inverse operators were applied on EEG data for calculation of minimum-norm estimates with a regularization parameter lambda2=1/9. Estimation of source activity and source-space statistical analysis was carried out using the MNE-Python (0.22.0) package (Gramfort et al., 2013). The estimated fixed-gaze and fixation-related source activities will be denoted by the FGSA and FRSA acronyms, respectively.

### 2.7 Statistical analyses

A lateralization index (LI) was calculated both for FGEA and FREA on trial level by subtracting the activity of left electrodes from the corresponding right ones. To control for the potential confounding effects of eye movements on FREA lateralization, eye-movement measures were regressed out from LI at subject level and trial filtering was also applied based on these variables beforehand (Supplementary Material, section 1.3). The average number of remaining trials was 869 (minimum: 414, maximum: 1625) for the natural reading experiment, while for the fixed-gaze paradigm about 130 trials left after elimination of artifactual data segments (W (average, minimum, maximum): 130.95, 106, 145; FF: 129.63, 101, 145; PR: 131.16, 102, 145). Individual-level average LI values entered group-level analyses. Single-trial LI of FGEA was averaged for each subject and condition (W, FF, PR), and group-level analysis of LI were run on these averages. Statistical analyses were carried out on a posterior cluster of occipital, occipito-temporal and parietal electrode pairs of special interest (PO8-PO7, PO4-PO3, P6-P5, O10-O9, PO10-PO9, O2-O1, P8-P7, P4-P3, P10-P9, PPO10h-PPO9h).

Group-level effects were assessed using the cluster-based permutation testing framework (Maris & Oostenveld, 2007; Nichols & Holmes, 2002) in the spatiotemporal domain (2D clustering) as well as using threshold-free cluster enhancement (TFCE; (Mensen & Khatami, 2013; Smith & Nichols, 2009)) with default settings (E=0.5, H=2, dh=0.1). Both methods were run with 1000 bootstrap repetitions using custom-written scripts, incorporating functions from the LIMO EEG toolbox (Pernet et al., 2011). For cluster-based permutation testing, the neighborhood matrix of EEG electrodes was generated by the ft_prepare_neighbours() function of the FieldTrip toolbox (Oostenveld et al., 2011) using the intersection of results obtained by triangulation and neighborhood distance methods. The threshold of neighborhood distance was set to 4.4 cm. Deviation of LI from zero was tested using the one-sample t-test. The effects of fixation rank on FREA and its lateralization were assessed by the paired samples t-test.

The relationship between the lateralization of the fixed-gaze N1 component and the lateralization of early FREA at the latency of first negative peaks was investigated using the skipped Spearman correlation analysis with 10000 bootstrap iterations as implemented in the Robust Correlation Toolbox (Pernet et al., 2013). Peak detection was performed using the average activity of left (P7, PO7, PO9, PPO9h) and right (P8, PO8, PO10, PPO10h) occipito-temporal electrode clusters. Details of peak detection are available in the Supplementary Material (section 1.4).

To assess the lateralization of fixed-gaze and fixation-related orthographic processing in source space, the activity of both hemispheres were morphed to the contralateral side in FreeSurfer’s fsaverage_sym space, an atlas with registration between the left and right hemispheres (Greve et al., 2013). In source space, hemispheric lateralization was characterized by the LIn normalized lateralization index. LIn was calculated for both FRSA and FGSA by subtracting the source activity of right vertices from the activity of corresponding left vertices and by dividing the difference with the sum of unsigned left and right source activities (Seghier, 2008). Accordingly, while the positive and negative sensor-space LI values denote more negative-going EEG deflection in left and right electrodes compared to the contralateral ones, respectively, the positive and negative source-space LIn values indicate stronger source activity at left-and right-hemispheric vertices compared to their contralateral pairs. The resulting LIn values were submitted to a two-tailed one-sample t-test with spatiotemporal clustering. The number of permutations was 1024, and the t statistic threshold was set to 2.026. The statistical significance of hemispheric lateralization was assessed for FRSA as well as for FGSA in conditions W and FF. Moreover, the difference of source-activity lateralization between the W and FF conditions was also tested by subtracting the lateralization in condition FF from the lateralization in condition W and submitting the resulting difference to a two-tailed one-sample t-test with spatiotemporal clustering. Permutation and t-threshold settings were the same as above. The obtained cluster P values (P_Clus_) were multiplied by two to correct for using two-tailed tests (P_corr_=P_Clus_×2), and clusters with P_corr_ values below 0.05 were considered significant. To identify brain regions exhibiting significant lateralization and to extract the average source activity of these regions of interest (ROI), cortical parcellation of the FSaverage anatomy was carried out by using the Human Connectome Project - Multi-modal Parcellation atlas (Glasser et al., 2016)) and a sub-parcellation of FreeSurfer’s 72 cortical regions (Fischl, 2004) into 448 labels as suggested in (Khan et al., 2018) and implemented in MNE-Python (Gramfort et al., 2013) as the aparc_sub atlas. ROIs were defined based on visual inspection of significant lateralization clusters shown on the left hemisphere. In order to extract the time course of source activity of left and the corresponding right ROIs, each participant’s cross-hemispheric data were morphed back to the FSaverage anatomy, for which the above atlases are available. The label of parcels used for definition of ROIs is provided in the Supplementary Material (section 1.5, Supplementary Table 1).

## 3 Results

### 3.1 Sensor-space results

The time course of grand-average FREA obtained during natural reading (Fig. 2a) was in agreement with previous findings and indicated a lateralization profile very similar to that we found recently for control participants without reading difficulties (Weiss, Nárai, et al., 2022). Statistical analysis of FREA lateralization was carried out in three time intervals defined by the zero-crossings of LI (94-170 ms, 170-236 ms and 236-310 ms) covering the period of early orthographic processing, similarly as in our recent study (Weiss, Nárai, et al., 2022). In the first time interval, a significant (P_Clust_=0.029) leftward lateralization of negative-going FREA deflection was revealed from 140 to 160 ms on posterior channel pairs. Considering the later time intervals, a significant (P_TFCE_<0.05, P_Clust_=0.001) rightward lateralization of negative-going FREA deflection was found from 175 to 225 ms and a significant (P_TFCE_<0.05, P_Clust_=0.001) rightward lateralization of positive-going FREA was revealed for the 245-295 ms time range in posterior electrodes.

**Figure 2.**
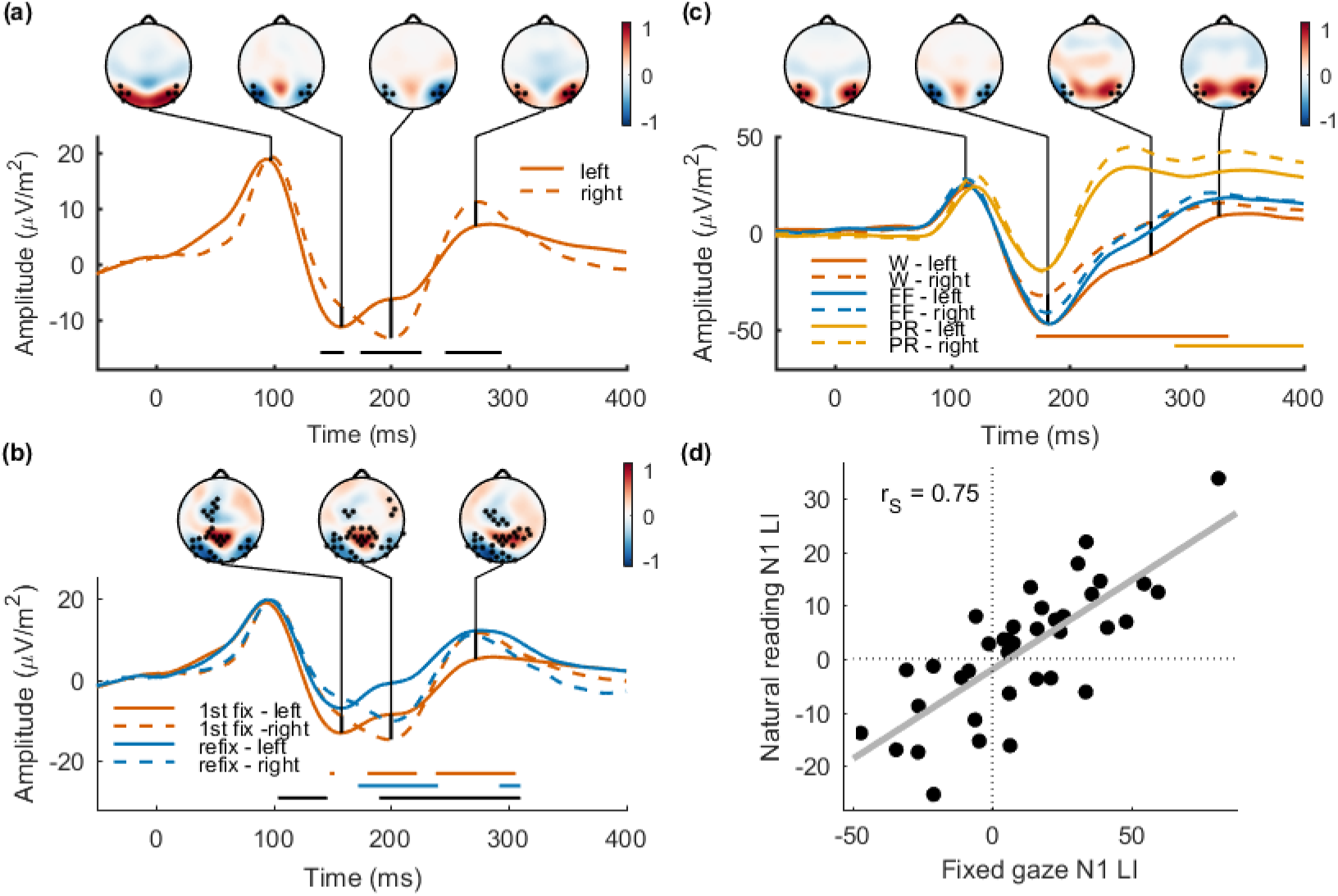
Lateralization of EEG activity in fixed-gaze and natural reading conditions. The time course of grand-average fixation-related EEG activity (FREA) recorded during natural reading is shown for left (PO9, PO7, P7, PPO9h) and right (PO10, PO8, P8, PPO10h) occipito-temporal electrode clusters (a). The location of the applied electrodes is indicated by black dots on the topographic plots showing normalized FREA at latencies of particular interest. Horizontal lines indicate significant lateralization of FREA. Grand-average FREA of first fixations and refixations (b) is presented for the same left and right occipito-temporal electrode clusters that were used in panel (a). Vermillion and blue horizontal lines denote significant lateralization of FREA for first fixations and refixations, respectively, and black horizontal lines indicate the significant difference between these lateralizations. Topographic plots represent the difference of FREA between first fixations and refixations. The black dots indicate the spatial distribution of significant fixation-rank effects at particular time points of special interest. Grand-average event-related EEG activity (FGEA) obtained for words (W), false-font strings (FF) and phase randomized noise stimuli (PR) during the fixed-gaze paradigm is provided in panel (c). The same left and right occipito-temporal electrode clusters were used as for FREA in (a). The location of applied electrodes is indicated by black dots on the topographic plots showing normalized FGEA for word stimuli at latencies of particular interest. Horizontal lines indicate significant lateralization of FGEA obtained for the W and PR conditions. The relationship between the occipito-temporal lateralization of FREA and FGEA at the latency of the first negative peaks is provided in panel (d). A significant positive correlation (r_S_=0.75, 95 % CI=[0.55 0.86]) was found between the EEG markers of early orthographic processing in fixed-gaze and natural reading conditions.

The effects of the fixation rank on FREA (Fig. 2b) were also very similar to the results obtained recently for control young adult readers (Weiss, Nárai, et al., 2022). Considering the occipito-temporal region, stronger negative deflection of FREA was found for the first fixations compared to the refixations predominantly in electrodes above the left hemisphere in the time range of early orthographic processing (125-400 ms, P_TFCE_<0.05, P_Clust_=0.001). This modulation of FREA by the rank of fixations resulted in significant differences of FREA lateralization between the first fixations and refixations (105-145 ms, P_TFCE_<0.05, P_Clust_=0.001; 190-235 ms, P_TFCE_<0.05, P_Clust_=0.001; 235-310 ms, P_TFCE_<0.05, P_Clust_=0.001). While for the first fixations significant FREA lateralization was found in all the three investigated time intervals (148-152 ms, P_TFCE_<0.05, P_Clust_=0.037 (140-155 ms); 180-222 ms, P_TFCE_<0.05, P_Clust_=0.001; 238-306 ms, P_TFCE_<0.05, P_Clust_=0.001), in the case of refixations significant FREA lateralization was found only for the second and third time ranges (172-236 ms, P_TFCE_<0.05, P_Clust_=0.001; 236-240 ms, P_TFCE_<0.05, P_Clust_=0.046; 292-310 ms, P_TFCE_<0.05, P_Clust_=0.018). The occipito-temporal effects of fixation rank on FREA were accompanied by centro-parietal and more anterior effects within the same significant cluster.

The time course of grand-average FGEA (Fig. 2c) indicates a modulation of occipito-temporal lateralization across the W, FF and PR conditions in the time range of the first negative FGEA component (N1). For word stimuli, significant (P_TFCE_<0.05, P_Clust_=0.001) leftward FGEA lateralization was revealed in posterior channel pairs from 175 to 335 ms, a time range that incorporates both the N1 and N2 FGEA components. Strings of false fonts generated a similar, but much weaker tendency of FGEA lateralization that did not reach the level of significance. Compared to the W and FF conditions, less prominent and bilateral deflection of FGEA was found for phase randomized noise stimuli at the latency of the N1 component. Moreover, no negative-going deflection of FGEA corresponding to the N2 component could be observed in the PR condition, and significant rightward lateralization of positive FGEA deflection was found starting only after the P2 component in the 290-400 ms time range (P_TFCE_<0.05, P_Clust_=0.002). Accordingly, the PR condition was excluded and only the W and FF conditions were considered during source-space analysis (section 3.2).

To investigate the relationship between the hemispheric lateralization of early orthographic processing in fixed-gaze and natural reading conditions, the lateralization index of occipito-temporal FGEA and FREA obtained at the latency of first negative components were correlated (Fig. 2d). A semiautomatic subject-level peak detection was performed on the left (PO9, PO7, P7, PPO9h) and right (PO10, PO8, P8, PPO10h) occipito-temporal channel-cluster averages. The automatic search for the individual peaks was started from the latency of the first negative left occipito-temporal grand-average peaks (158 ms for FREA and 182 ms for FGEA in condition W). Subject-level occipito-temporal LI was averaged within a 10 ms (5 sample points) long time window centered at the latency of the earlier negative peak regardless of whether this peak occurred in the left or right occipito-temporal region. Accordingly, these average LIs do not reflect the relationship of left and right N1 peak amplitudes after controlling for their difference in latency, but they simply reflect the difference between the amplitude of the earlier negative peak and the corresponding FGEA/FREA deflection on the contralateral side. Using these lateralization measures calculated for FGEA and FREA, a significant positive correlation (r_S_=0.75, 95 % confidence interval CI=[0.55 0.86]) was found between the lateralization of EEG activity in fixed-gaze and natural reading conditions. The positive correlation indicates a high percentage of participants (81.6 %) with left- (Fig. 2d, upper right quadrant) or right-lateralized (Fig. 2d, lower left quadrant) EEG activity in both fixed-gaze and natural reading conditions at the latency of the first negative EEG peaks.

### 3.2 Source-space results

To investigate the brain sources contributing to the lateralization of sensor-space EEG correlates of orthographic processing in fixed-gaze and natural reading conditions, distributed source estimates of FGEA and FREA were calculated. Significant lateralization of fixation-related source activity (FRSA) was found in all the three time intervals (140-160 ms, 175-225 ms, 245-295 ms) indicated by sensor-space analysis (Fig. 3a). Two significant positive clusters were obtained in the time range of early orthographic processing between 140 and 160 ms (Fig. 3a, top row). A posterior significant cluster (P_corr_=0.012) indicated stronger FRSA in left compared to right lateraloccipital (LO), inferior parietal (IPT) and superior parietal (SPT) brain regions. Left-hemisphere dominance of FRSA was found also for superior premotor (SPM) and inferior premotor (IPM) cortical areas reflected by a significant frontal cluster (P_corr_=0.032). The time course of grand-average FRSA in ROIs covering the brain regions with significant lateralization (Fig. 3b) indicates higher FRSA in the left compared to the right IPM area, starting from about 100 ms (the latency of the FREA lambda peak) and lasting up to about 200 ms. The lateralization of FRSA in the IPM area is accompanied by stronger FRSA in left compared to right SPM, SPT, IPT as well as LO areas around 150 ms. The strength and the duration of these hemispheric differences varied across the regions. The stronger FRSA in left posterior brain regions most likely reflects the stronger negative-going deflection of left compared to right occipito-temporal FREA in this time range. No lateralization of FRSA could be observed in anterior (AV) and middle ventral temporal (MV) areas in this time interval. Interestingly, similarly to the LO area, a left-hemisphere dominance of mean FRSA can be seen for the posterior ventral occipito-temporal (PV) ROI, however, this lateralization did not reach the level of significance (see Fig. 3a, top row, posterior ventral region), most likely due to stronger attenuation of source activity in left compared to right ventral occipito-temporal areas showing also a posterior to anterior gradient (Fig. 3b, right column). In the 175-225 ms time range (Fig. 3a, middle row), a single significant cluster (P_corr_=0.002) was found. This negative cluster indicates a rightward lateralization of FRSA predominantly in the MV temporal cortical areas corresponding to the stronger negative deflection of FREA in right compared to left posterior EEG channels in this time interval. Similarly, a single significant negative cluster (P_corr_=0.014) was found also for the 245-295 ms time range (Fig. 3a, bottom row) reflecting stronger FRSA in right compared to left anterior and middle ventral cortical areas. This effect corresponds to the stronger positive deflection of right compared to left occipito-temporal FREA found for this time interval by sensor-space lateralization analysis. Visual inspection of mean FRSA also indicated a stronger activity in left compared to right SPM and IPM brain regions around 200 ms. This tendency was supported by a marginally significant positive cluster (P_Clus_=0.07) suggesting left-lateralized FRSA not only for the SPM and IPM ROIs, but also for cortical areas anterior and superior to SPM (Supplementary Fig. 1).

**Figure 3.**
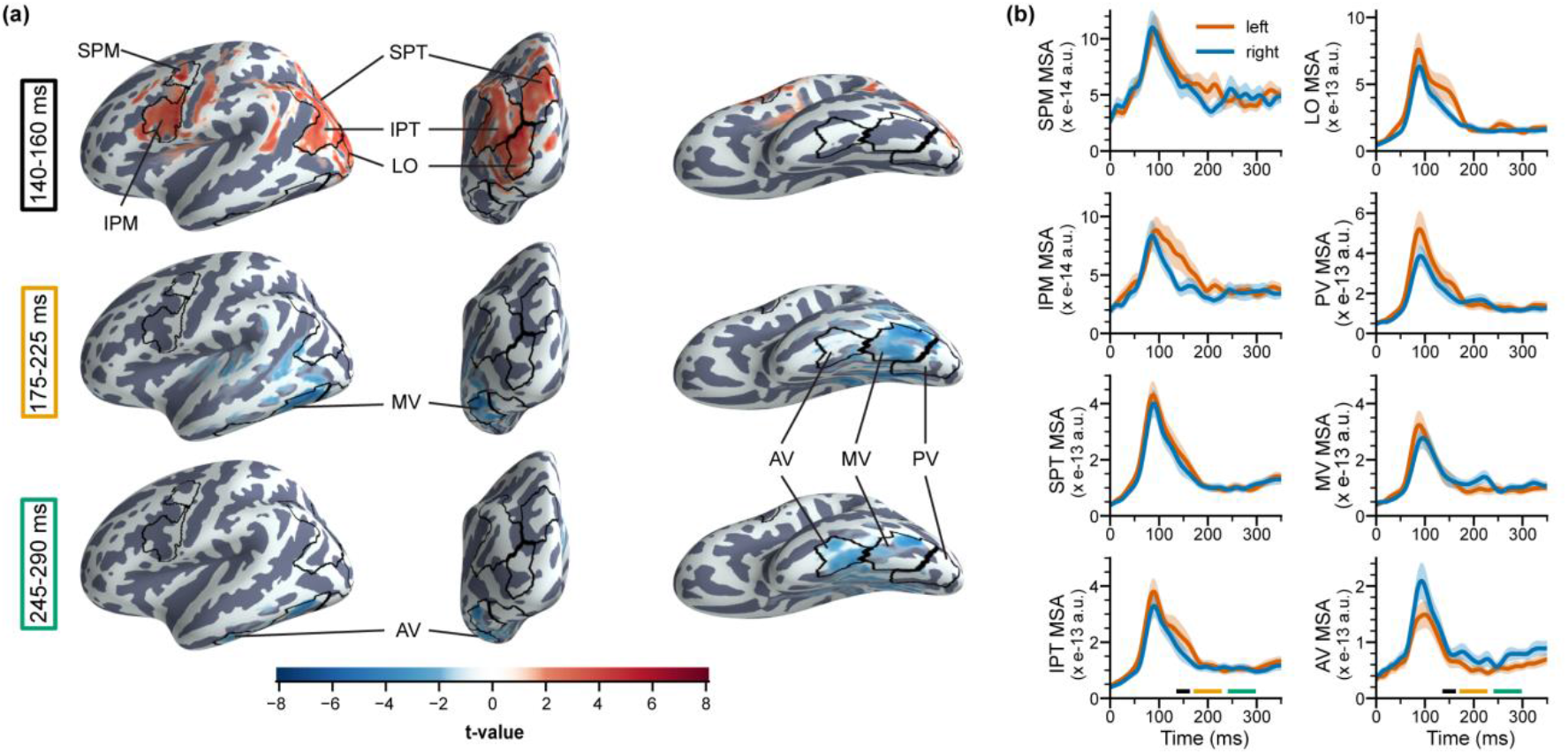
Lateralization of fixation-related source activity. Panel (a) shows the topography of averaged t-values that correspond to significant spatiotemporal samples of the lateralization clusters (P_corr_<0.05) obtained for fixation-related source activity (FRSA) in the 140-160 ms (top), 175-225 ms (middle) and 245-295 ms (bottom) time intervals. Mean source activity (MSA) of brain regions of special interest (ROIs) is shown both for left and right hemispheres (b). The time course of MSA was obtained by averaging the FRSA of vertices belonging to particular ROIs. Shaded areas correspond to standard error of mean. The horizontal solid lines denote the time intervals used for both statistical analyses and averaging of t-values in (a). The boundary of the ROIs is denoted by black lines on the FSaverage anatomy of the left hemisphere using lateral (left column), caudal (middle) and ventral (right) views in panel (a). Abbreviations: LO: lateraloccipital; SPT: superior parietal; IPT: inferior parietal; SPM: superior premotor; IPM: inferior premotor; AV: anterior ventral temporal; MV: middle ventral temporal; PV: posterior ventral occipito-temporal.

Contrasting the lateralization of fixed-gaze source activity (FGSA) obtained for word and false-font stimuli, a single significant cluster (P_corr_=0.002) was found in the 175-335 ms time range indicating significant effects in LO, MV and PV areas as well as at the temporo-parieto-occipital (TPO) junction (Fig. 4a). The time course of mean lateralization in these ROIs (Fig. 4b) shows more positive-going lateralization indices for words compared to false fonts within the time interval of statistical analysis suggesting stronger left-hemispheric lateralization for words that is in agreement with lateralization results obtained for W and FF conditions in sensor space (Fig. 2c). Furthermore, two peaks could be observed in these mean lateralization time courses within the 175-220 ms and 220-290 ms time intervals, most likely corresponding to the N1 and N2 FGEA components in sensor space.

**Figure 4.**
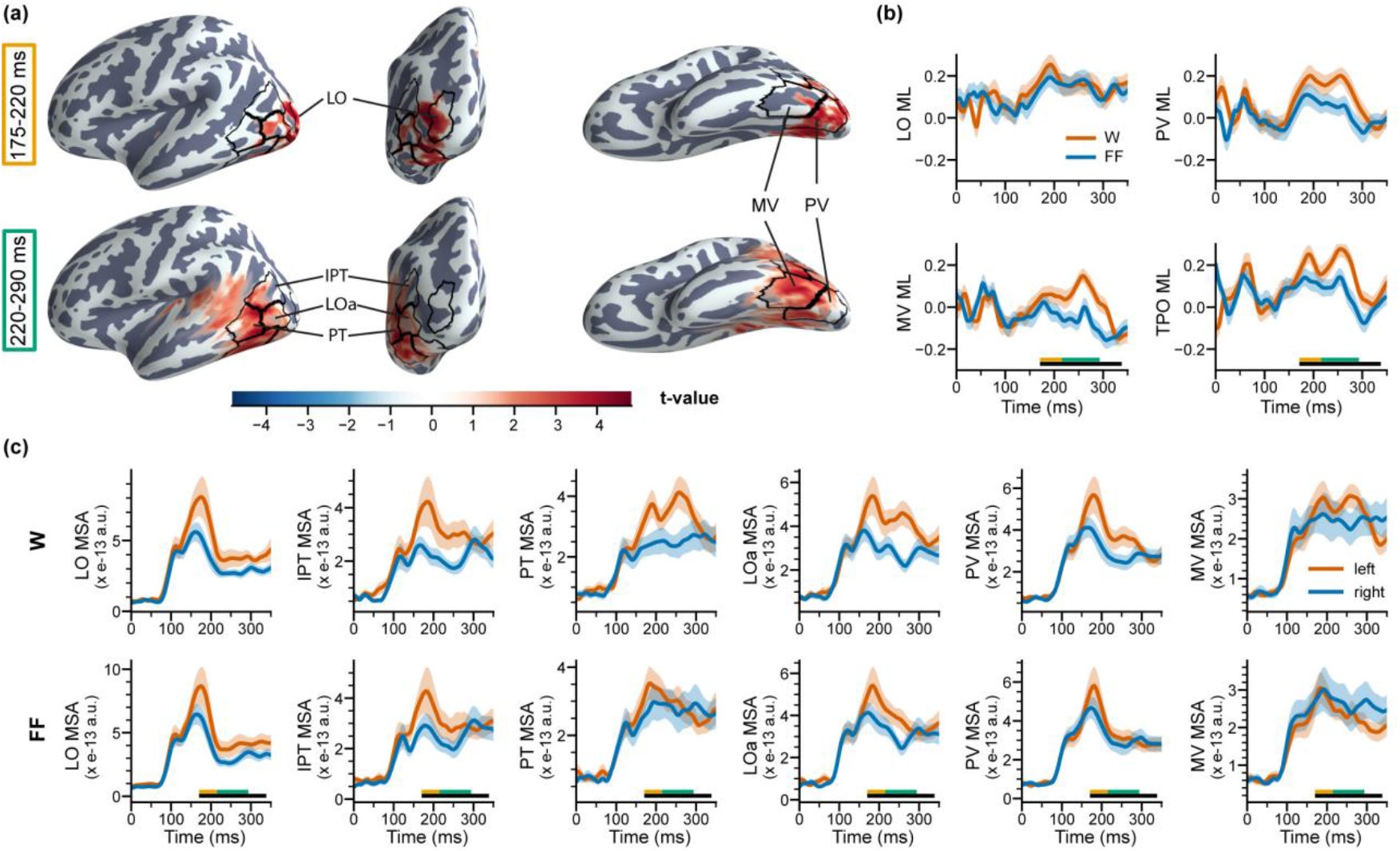
The difference between the lateralization of fixed-gaze source activity during processing of words and false-font stimuli. Panel (a) shows the topography of averaged t-values that correspond to the significant cluster (P_corr_=0.002) obtained for testing the difference of fixed-gaze source activity (FGSA) lateralization between the W (words) and FF (false fonts) conditions. Statistical analysis was performed for the 175-335 ms time interval indicated by the sensor-space results. However, averaging of t-values was carried out by considering the significant spatiotemporal samples in the 175-220 ms (top) and 220-290 ms (bottom) time intervals separately as suggested by the time course of grand-average lateralization indices (b) and grand-average FGSA (c) shown for brain regions of special interest (ROIs). Mean lateralization (ML) of FGSA is provided for the W and FF conditions (b), while mean source activity (MSA) is presented for the two conditions and both for the left and the corresponding right ROIs (c). The time course of ML and MSA was obtained by averaging the lateralization indices and the FGSA of vertices belonging to the ROIs, respectively. Shaded areas correspond to standard error of mean. The horizontal black lines denote the time interval used for statistical analysis, while the colored horizontal lines indicate the time intervals used for averaging of t-values in (a). The boundary of the ROIs is denoted by black lines on the FSaverage anatomy of the left hemisphere using lateral (left), caudal (middle) and ventral (right column) views in panel (a). The topography of significant lateralization clusters obtained for the W and FF conditions is shown in Fig. 5. Abbreviations: LO: lateraloccipital; LOa: anterior lateraloccipital; IPT: inferior parietal; PT: posterior temporal; TPO: temporo-parieto-occipital, combination of the PT, IPT and LOa ROIs; MV: middle ventral temporal; PV: posterior ventral occipito-temporal.

The topography of average t-values indicates condition effects on FGSA lateralization predominantly in LO, anterior LO (LOa) and PV ROIs from 175 to 220 ms (Fig. 4a, top) and in more anterior MV and posterior temporal (PT) areas between 220 and 290 ms (Fig. 4a, bottom). The transition from earlier posterior to later anterior effects can be understood by inspecting the mean FGSA of left and right ROIs in both conditions (Fig. 4c). While the earlier condition effects seem to be driven by the left-lateralized early FGSA in posterior ROIs in both conditions, the later condition effects might originate from left-lateralized FGSA present in more anterior MV and PT cortical areas observed for word stimuli only. We assume that the earlier FGSA peaks in posterior LO and PV areas correspond to processing of familiar letter (W) and unfamiliar letter-like (FF) visual stimuli, while the later peak observed for words only probably reflects higher-level lexical processing required by word stimuli only. Accordingly, the significantly stronger leftward lateralization of FGSA found for words compared to false fonts in LO and PV areas in the 175-220 ms time range might indicate expertise effects, enhanced sensitivity for familiar letters compared to unfamiliar, but still letter-like stimuli (false fonts).The topography of significant lateralization clusters obtained for word (P_corr_=0.002, Supplementary Fig. 2a) and false-font (P_corr_=0.038, Supplementary Fig. 2b) stimuli supports these assumptions by showing significant left-hemispheric lateralization of FGSA in posterior LO and PV areas for both types of stimuli in the earlier time range and in medial and posterior ventral areas predominantly for word stimuli in the later time interval. The higher t-values obtained for words compared to false-font stimuli confirm the significantly stronger lateralization of FGSA presented for words in Fig. 4. Finally, although a stronger left-hemispheric FGSA lateralization was found for words compared to false fonts also in the TPO region indicating stronger effects especially in the PT and LOa cortices from 220 ms to 290 ms (Fig. 4), the significant lateralization obtained for both W and FF conditions in this region (Supplementary Fig. 2) might also reflect processes such as visual short-term memory required by the one-back task in both conditions (Benedictis et al., 2014).

## 4 Discussion

Lateralization of orthographic processing has been shown to reflect reading expertise and indicate impaired reading. However, despite its general importance in characterizing reading processes, research of orthographic processing lateralization was limited to fixed-gaze experimental paradigms until recently. Accordingly, our knowledge on how earlier findings obtained in fixed-gaze experimental conditions translate to natural reading is still very limited. To address this shortcoming, here we evaluated the lateralization of orthographic processing in fixed-gaze and natural reading conditions within the same participants without reading difficulties. Markers of hemispheric lateralization were assessed by both sensor-space and source-space analyses.

In sensor-space, we validated earlier findings on lateralization of fixed-gaze and fixation-related EEG activity recorded during orthographic processing. For the fixed-gaze EEG activity, we found left-lateralized negative-going deflection in occipito-temporal electrodes during processing of word stimuli. In line with previous studies, two negative components could be identified on grand-average EEG activity within the time interval of significant effects. The first negative component with a peak latency at around 180 ms most likely corresponds to letter-level extraction of orthographic information, while the second negative component in the 250-300 ms time range presumably reflects word-level processing (Bentin et al., 1999; Cohen et al., 2000; Rossion et al., 2003). False-font stimuli evoked similar EEG activity as word stimuli. However, the tendency of left-lateralized negative deflection did not reach the level of significance for false fonts in sensor-space in line with earlier findings indicating left lateralized and rather bilateral N1 ERP components for familiar and unfamiliar orthographic stimuli, respectively (Maurer et al., 2008). These results support the importance of reading acquisition and reading experience in shaping the lateralization of orthographic processing (Brem et al., 2010). Considering the lateralization of fixation-related EEG activity, we confirmed our recent findings obtained for control participants during natural reading of sentences presented with default inter-letter spacing (Weiss, Nárai, et al., 2022). We obtained significant left and then right lateralized FREA corresponding to the first negative and latter components, respectively. Moreover, we also validated the dependence of FREA and its lateralization on the rank of fixations. Our results confirm that early orthographic processing at the latency of the first negative FREA component is lateralized only in the case of first fixations. To address the need for relating the EEG correlates of fixed-gaze and natural reading (Kornrumpf et al., 2016; Niefind & Dimigen, 2016; Weiss et al., 2016), we assessed the correlation between the early occipito-temporal lateralization of EEG activity in fixed-gaze and natural reading conditions. Importantly, the positive relationship found between the lateralization of fixed-gaze and fixation-related EEG activity at the latency of the first negative peaks suggests common neural processes contributing to early letter-level orthographic processing in both reading conditions. Although the experimental paradigms applied in this study allowed the validation of most elementary findings on the lateralization of orthographic processing in both fixed-gaze and natural reading conditions, they are limited in providing a fine-grained assessment of fixed-gaze lateralization markers and replication of letter spacing and reading ability effects on the lateralization of fixation-related EEG activity. More detailed information could be obtained on the lateralization of brain activity at different levels of orthographic processing by also considering consonant string, nonword, pseudoword as well as high- and low-frequency word stimuli.

The latency of the N1 component (∼180 ms) in our fixed-gaze paradigm is somewhat larger than one might expect for word stimuli, and this may also contribute to the later onset of lateralization effects in fixed-gaze compared to natural reading mode. Although the first negative occipito-temporal EEG component evoked by visual word stimuli in fixed-gaze conditions is often peaking as early as 150 ms after stimulus onset (Grainger & Holcomb, 2009), the latency of the N1 component is highly variable. It depends on multiple stimulus features such as contrast, stimulus extent and eccentricity (Busch et al., 2004; Schindler et al., 2018). As it has been shown that decreasing stimulus size increases the latency of the P1 and N1 components (Busch et al., 2004; Schindler et al., 2018), the delay in the latency of the fixed-gaze N1 component could be explained by the relatively small font size and shorter words compared to natural reading stimuli that resulted in fixed-gaze stimulus extent below two visual degrees in this study.

To reveal the lateralization correlates of orthographic processing at the level of brain sources, we estimated distributed fixation-related source activity. In the first time interval of FREA lateralization, around 150 ms after the onset of fixations, stronger fixation-related source activity was found in left compared to right LO, IPT, SPT, IPM and SPM brain regions. The inferior and superior parietal areas along with the inferior and superior premotor regions are known to form the fronto-parietal attentional network, subserving oculomotor behavior and relocation of visuospatial covert attention (Behrmann et al., 2004; Bisley & Goldberg, 2010; Buschman & Miller, 2007; Colby & Goldberg, 1999; Corbetta & Shulman, 2002; Kastner et al., 2007; Silver & Kastner, 2009), two inherent processes of natural reading. In agreement with our results, directing the visuospatial covert attention to the right visual field as well as preparation and execution of left-to-right saccadic eye movements have been associated with leftward lateralization of brain activity in these areas (Corbetta & Shulman, 2002; Kastner et al., 1999). Shifts of covert attention might contribute to presaccadic preview, while initiation and execution of following saccades could be reflected by the marginally significant lateralization found in IPM, SPM and supplementary motor areas around 200 ms. However, as visuospatial attention and eye movements are not segregated processes (Corbetta, 1998), their dissociation is not feasible based on our data. No lateralization of source activity was found in these brain regions for fixed-gaze orthographic processing, supporting the assumption that the above effects obtained for fixation-related source activity reflect neural processes specific for natural reading. According to these results, the impaired posterior lateralization of early orthographic processing during natural reading in dyslexic readers (Weiss, Nárai, et al., 2022) might also support recent findings suggesting visuospatial attention deficits in dyslexics (Collis et al., 2013; Vidyasagar, 2019; Vidyasagar & Pammer, 2010; White et al., 2019). However, this assumption has to be validated by further studies applying source-space analysis of fixation-related brain activity during natural reading in dyslexic and control participants.

The early lateralization in fronto-parietal areas was accompanied by leftward lateralization of source activity in the lateraloccipital region. Despite their moderate selectivity to word stimuli (Kadipasaoglu et al., 2016), the importance of extrastriate occipital areas in processing of orthographic information is supported by multiple lines of experimental evidence (Barzegaran & Norcia, 2020; Boros et al., 2016; Gold & Rastle, 2007; Hervais-Adelman et al., 2019; Posner et al., 1988; Puce et al., 1996; Temple et al., 2001; Woolnough et al., 2020). These studies suggest left-lateralized sensitivity to morphological features of visual word stimuli and specialization of extrastriate occipital areas for an intermediate processing of perceptual features of letter strings between early striate and later lexico-semantic processing in more anterior cortical areas along the ventral occipito-temporal stream. In two fMRI studies (Boros et al., 2016; Temple et al., 2001), using tasks targeting orthographic processing, dyslexic children showed a significant underactivation in the left middle occipital gyrus (Temple et al., 2001) with a peak group difference obtained for letter stimuli in the left lateral occipital cortex (Boros et al., 2016). These findings seem to be in agreement with our recent results showing impaired lateralization of early orthographic processing during natural reading in dyslexic readers (Weiss, Nárai, et al., 2022) that might also indicate visuospatial attention deficits in dyslexics (Collis et al., 2013; Vidyasagar, 2019; Vidyasagar & Pammer, 2010; White et al., 2019). In the present study, the lateralization of fixed-gaze source activity from 170 to 220 ms confirmed the contribution of the lateraloccipital area to early orthographic processing of words as well as false-font stimuli without meaning, indicating a general sensitivity of this region to letter-like patterns. Moreover, the significantly stronger lateralization obtained for words compared to false fonts might indicate beneficial effects of reading expertise on efficiency of orthographic processing (Brem et al., 2010; Maurer et al., 2008).

The early fixed-gaze lateralization in the LO region was accompanied by significant lateralization in posterior ventral cortical areas also shown to be involved in early letter-level processing (Thesen et al., 2012). In the case of natural reading, the same tendency of lateralization was observed for the posterior ventral areas as for the lateraloccipital region. However, the lateralization in the posterior ventral region did not reach the level of significance, most likely due to stronger suppression of brain activity along the left ventral occipito-temporal stream (see Fig. 3b, right column) that might also correspond to repetition priming (Dimigen et al., 2012; Temereanca et al., 2012) in the form of FREA modulation by presaccadic preview (Buonocore et al., 2020; Kornrumpf et al., 2016). The posterior-anterior gradient of this suppression in the left occipito-temporal areas resulted in a significant rightward lateralization of fixation-related source activity predominantly in middle ventral cortical areas from 175 to 225 ms as well as in middle and more anterior ventral cortical areas in the 245-290 ms time interval. These effects most likely reflect word-form identification in the VWFA and higher-level lexico-semantic processing in the VWFA and more anterior inferotemporal areas (Halgren et al., 1994, 2006; Temereanca et al., 2012) in agreement with recent findings indicating sequential and then interactive processing of letter and word stimuli (Lerma-Usabiaga et al., 2018; Thesen et al., 2012). The lateralization of fixed-gaze source activity found for words, but not for false-font stimuli in the medial ventral temporal region from 220 to 290 ms supports the assumption of lexical processing in the medial ventral ROI containing the VWFA, and also suggests earlier processing of lexical content during natural reading due to preview and anticipatory processes. Finally, the lack of lateralization in anterior ventral areas during fixed-gaze word processing without a need for extraction of semantic content might also confirm the assumption of higher-level processing in the 220-290 ms time interval during natural reading of sentences that definitely requires extraction of semantic content for integration of contextual information across multiple words.

We acknowledge that as the solution of the inverse problem used for estimation of source activity is undetermined, analysis parameters, such as the settings of the forward model and the inverse method might influence the resulting reconstructed source estimates (Akalin Acar & Makeig, 2013; Mahjoory et al., 2017). Usage of template anatomy instead of individual anatomical data may result in less accurate source estimates, especially if the number of sensors is low (Akalin Acar & Makeig, 2013). Although using a template anatomy with a high-density 96-channel electrode system we obtained source-space results that were in agreement with earlier findings and showed consistency across the experimental paradigms, further studies employing localizer paradigms combined with individual MRI anatomies are needed to verify our findings.

We designed the fixed-gaze experiment with the intention to replicate traditional ERP studies so we can interpret our results in the context of the extensive literature on word-evoked EEG activity. One limitation of this approach is the inevitable difference in gaze duration and stimulus set between the two experimental paradigms. Accordingly, performing studies with matched words and gaze duration can be an important next step in investigating the underlying processes of the obtained EEG lateralization profiles in fixed-gaze and natural reading modes. However, while using the same words in the fixed-gaze paradigm as in the sentences used for natural reading seems to be straightforward, matching gaze duration can be challenging. Gaze duration is not only dependent on word length and frequency (Kliegl et al., 2004; Loberg et al., 2019) but also on parafoveal preview (Kornrumpf et al., 2016). As the benefit of parafoveal preview interacts with word length in natural reading and parafoveal preview is not available in standard fixed-gaze paradigms applying one-by-one presentation of orthographic stimuli, more sophisticated fixed-gaze paradigms applying also flanker words and shifted presentation of stimuli might be needed for better matching of fixed-gaze and natural reading processes (Kornrumpf et al., 2016). The interaction between word length and parafoveal preview can be avoided by limiting the length of words to a narrow range. Although such limitation of word length might prevent the usage of connected text stimuli, reading of unconnected words in saccadic and fixed-gaze modes could still aid the identification of effects of parafoveal preview and eye movements on lateralization of orthographic processing. Moreover, as parafoveal preview might rely on covert shifts of visuospatial attention the direction of which depends on the direction of reading, the effects of parafoveal preview and eye movements on lateralization of orthographic processing could also be investigated by performing natural reading experiments in languages applying different reading directions. Since previous studies using fixed-gaze experimental paradigms have shown that the leftward lateralization of the N1 ERP component is not limited to alphabetic languages but also generalizes to logographic scripts (Maurer et al., 2008; Yum et al., 2011), the effects of parafoveal preview and eye movements on lateralization of orthographic processing could also be assessed in different reading directions while using the same logographic language.

## 5 Conclusions

Here we confirmed the leftward lateralization of fixed-gaze orthographic processing and also validated that early leftward lateralization of fixation-related EEG activity is followed by rightward lateralization during natural reading. Importantly, we found a strong positive relationship between the early leftward lateralization in fixed-gaze and natural reading conditions indicating common neural mechanisms subserving early orthographic processing. For this time range, we revealed left-lateralized brain activity in lateraloccipital and posterior ventral occipito-temporal regions most likely corresponding to letter-level processing. In the subsequent time intervals, we obtained left-lateralized medial ventral temporal and right-lateralized medial and anterior ventral temporal brain activity for fixed-gaze and natural reading conditions, respectively, suggesting suppression of left ventral temporal cortical activity at these later stages of word- and presumably higher-level processing of orthographic information during natural reading. Accordingly, our results corroborate earlier findings and provide deeper insight into lateralization of orthographic processing during natural reading. The results demonstrate that although some findings obtained for fixed-gaze processing of orthographic information translate nicely to reading with saccadic eye movements, reading research applying ecologically valid conditions is essential to unveil the brain mechanisms subserving attentional, oculomotor and higher-level processes specific for natural reading.

## Acknowledgments

This work was supported by the Hungarian Brain Research Program 2.0 [NAP 2.0 4001-17919]; and the Hungarian Scientific Research Fund [K112093]. The authors thank Dávid Farkas and Dr. Dénes Tóth for their contribution to the preparation of text stimuli. The assistance of Dr. Nándor Donauer in applying for ethical approval is highly appreciated. The authors are also grateful to Dr. Marcus Nyström for sharing the implementation of the adaptive eye-tracking data analysis algorithm.

