## Supplementary Material for "Lateralization of orthographic processing in fixed-gaze and natural reading conditions"

^*^ Corresponding author

1117 Budapest

Hungary

### 1 Supplementary methods

#### 1.1 Validation of eye tracking

Quality of eye tracking was validated before the presentation and after reading of all sentences by asking the participants to gaze at fixation points presented to the left and to the right of text stimuli, respectively. To initiate the presentation of sentences, gaze position had to remain within a 1°×1° validation square around the left fixation point for at least 150 ms. The validation timeout was 3 s. If the ET validation failed before the presentation of the text lines, the ET device was recalibrated. A very similar validation procedure was applied at the end of text lines using the fixation points presented right to the end of text stimuli. However, on the right side of texts, the validation criterion was relaxed if necessary, and the initial validation time was only 2 s. The relaxed validation criterion required the horizontal gaze position to be right to a vertical margin assumed at the horizontal position of the right fixation points minus 1/9° for 150 ms at text line endings. The experimenter had the opportunity to manually skip the gaze validation procedure at the end of text lines if the relaxed criterion was not met. Manual skipping was marked to allow offline exclusion of sentences with low ET accuracy from further analyses.

#### 1.2 Timing of trigger signals and messages

Timing of trigger signals and messages sent to the EEG and ET devices was assessed by multiple tests. In the case of EEG, timing of trigger signals was compared to the latency of responses recorded by a photodiode, while the synchrony of messages sent to the two devices was investigated by comparing the inter-message intervals in the two datasets and by inspecting the latency of fixation-related EEG components. No significant lags and jittering was found in the timing of trigger signals and messages, the latency of fixation-related EEG components reflecting eye-movement artefacts and early visual responses was in agreement with the assumptions and earlier findings published by different research groups (Dimigen et al., 2011; Plöchl et al., 2012; Weiss et al., 2016). The corneo-retinal and spike artefacts related to eye movements were observed around the latency of saccade end times used for triggering of FREA. These artefacts appeared on raw subject-level FREA averages as well as in the activity of corresponding independent components. Moreover, the latency of the first positive occipital and occipito-temporal FREA peaks was around 100 ms (Fig. 2a) in line with previous findings.

#### 1.3 Usage of eye-movement measures during EEG analyses

Seven eye-movement measures, the amplitude of incoming, preceding and outgoing saccades, the duration of current, previous and following fixations and the horizontal gaze position of current fixations were from lateralization indices. FREA trials were filtered based on ET features. Trials with any of the seven ET covariates missing, those with regressive incoming saccade or incoming saccade amplitude below 0.26 or above 10 visual degrees were excluded. Furthermore, only trials with a horizontal position of current fixation falling on some of the words were used. Preceding and outgoing saccades with more than one second between their and incoming saccade endings were regarded as missing. Previous fixations ending before the beginning of the incoming saccade with more than 0.1 seconds were excluded. Similarly, current and following fixations were excluded when their beginning followed the end of the preceding saccade with more than 0.1 seconds.

#### 1.4 Peak detection

Peak detection was performed using the average activity of left (P7, PO7, PO9, PPO9h) and right (P8, PO8, PO10, PPO10h) occipito-temporal electrode clusters. The detection of subject-level N1 FGEA peaks started within an 80 ms long time window centered on the latency of the grand-average left occipito-temporal FGEA peak. If local minimum was not found in this time range, the length of the window was increased gradually and symmetrically, up to 200 ms. In the case of natural reading, two early negative FREA peaks could be observed after the lambda peak within the 100-200 ms time window in several subjects. To differentiate between the two peaks, we applied an asymmetrical peak search. The detection of the earlier negative FREA peaks used in correlation analysis was started in a 20 ms time window ending at the latency of the left grand-average occipito-temporal FREA peak. The length of the window increased gradually and symmetrically up to 200 ms until a local minimum was found. Peak detection was validated by visual inspection and manual correction was applied for three subjects.

#### 1.5 Regions of interest

**Supplementary Table 1 Regions of interest.** The list of parcel labels is provided for all regions of interest (ROIs). Cortical parcellation of the FreeSurfer’s FSaverage anatomy (Fischl, 2012) was carried out by using the Human Connectome Project - Multi-modal Parcellation atlas (HCP-MMP, (Glasser et al., 2016)) and a sub-parcellation of FreeSurfer’s 72 cortical regions (Fischl, 2004) into 448 labels as suggested in (Khan et al., 2018) and implemented in MNE-Python (Gramfort et al., 2013) as the aparc_sub atlas.

| **ROI** | **Label 1** | **Label 2** | **Label 3** | **Label 4** | **Label 5** | **Label 6** | **Atlas** |
| --- | --- | --- | --- | --- | --- | --- | --- |
| SPM | L_FEF_ROI-lh | L_55b_ROI-lh |  |  |  |  | HCP-MMP |
| IPM | L_6r_ROI-lh | L_6v_ROI-lh | L_IFJa_ROI-lh | L_IFJp_ROI-lh |  |  |  |
| IPT | inferiorparietal_1-lh | inferiorparietal_2-lh | inferiorparietal_3-lh | inferiorparietal_4-lh | inferiorparietal_7-lh | inferiorparietal_9-lh | aparc_sub |
| SPT | superiorparietal_10-lh | superiorparietal_11-lh | superiorparietal_12-lh | superiorparietal_13-lh |  |  |  |
| AV | inferiortemporal_3-lh | inferiortemporal_4-lh | inferiortemporal_6-lh - anterior section |  |  |  |  |
| MV | lateraloccipital_9-lh - ventral section | inferiortemporal_6-lh - posterior section | inferiortemporal_7-lh - ventral section | inferiortemporal_8-lh - ventral section | fusiform_3-lh | fusiform_4-lh |  |
| PV | lateraloccipital_10-lh | lateraloccipital_11-lh | fusiform_1-lh | fusiform_2-lh |  |  |  |
| LO | lateraloccipital_2-lh | lateraloccipital_3-lh |  |  |  |  |  |
| IPT (TPO) | inferiorparietal_3-lh | inferiorparietal_4-lh |  |  |  |  |  |
| PT (TPO) | inferiortemporal_8-lh - dorsal section | middletemporal_1-lh | middletemporal_2-lh |  |  |  |  |
| LOa (TPO) | lateraloccipital_7-lh - anterior section | lateraloccipital_8-lh | lateraloccipital_9-lh - dorsal section |  |  |  |  |

Abbreviations: SPM: superior premotor; IPM: inferior premotor; IPT: inferior parietal; SPT: superior parietal; AV: anterior ventral temporal; MV: middle ventral temporal; PV: posterior ventral occipito-temporal; LO: lateraloccipital; IPT (TPO): inferior parietal as a part of the TPO ROI; PT (TPO): posterior temporal; LOa (TPO): anterior lateraloccipital; TPO: temporo-parieto-occipital, combination of the IPT (TPO), PT (TPO) and Loa (TPO) ROIs.

### 2 Supplementary results

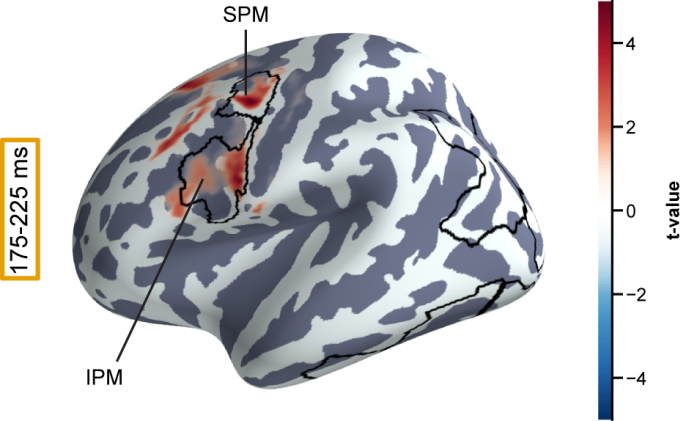

**Supplementary Figure 1 The topography of the marginally significant frontal cluster obtained by lateralization analysis of fixation-related source activity in the 175-225 ms time range.** The presented topography reflects the average of t-values that correspond to significant spatiotemporal samples belonging to the marginally significant cluster (P_Clus_=0.07). The mean source activity of indicated brain regions of special interest (ROI) is shown in Fig. 3b. The ROIs shown in this figure correspond to those used in Fig. 3a. The boundary of the ROIs is denoted by black lines on the FSaverage anatomy of the left hemisphere using the lateral view. Abbreviations: SPM: superior premotor; IPM: inferior premotor.

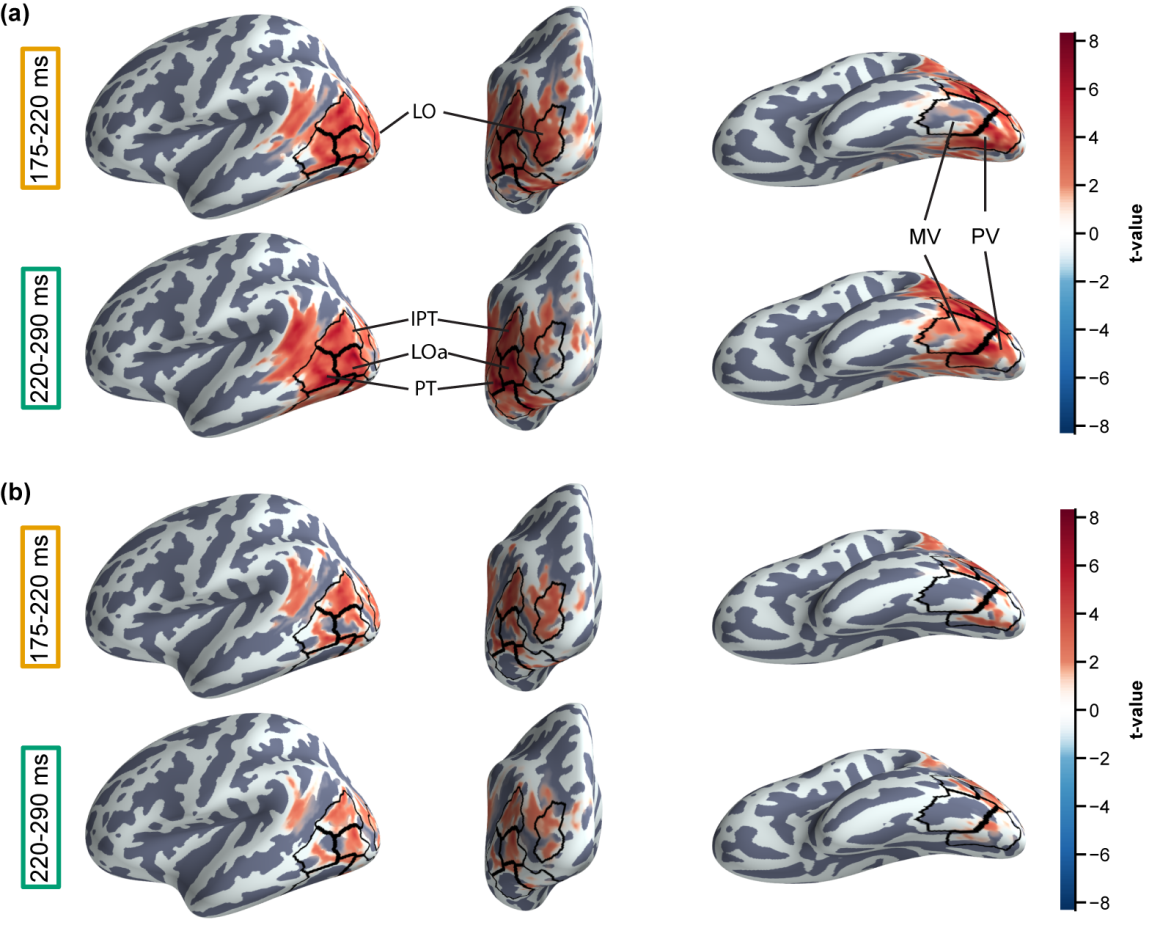

**Supplementary Figure 2 The topography of fixed-gaze source-activity lateralization during processing of words and false-font stimuli.** The topographies of averaged t-values that correspond to the significant lateralization clusters obtained for fixed-gaze source activity (FGSA) in conditions W (words, Pcorr=0.002) and FF (false fonts, Pcorr=0.038) are shown in panels (a) and (b), respectively. Statistical analyses were performed for the 175-335 ms time interval indicated by the sensor-space results. However, averaging of t-values was carried out by considering the significant spatiotemporal samples in the 175-220 ms (top) and 220-290 ms (bottom) time intervals separately as suggested by the time course of grand-average FGSA and lateralization shown for brain regions of special interest (ROIs) in Fig. 4, panels (b) and (c). The ROIs shown in this figure correspond to those used in Fig. 4a. The boundary of the ROIs is denoted by black lines on the FSaverage anatomy of the left hemisphere using lateral (left column), caudal (middle) and ventral (right) views. Abbreviations: LO: lateraloccipital; LOa: anterior lateraloccipital; IPT: inferior parietal; PT: posterior temporal; MV: middle ventral temporal; PV: posterior ventral occipito-temporal.
